# Hypoxia-mediated regulation of DDX5 through decreased chromatin accessibility and post-translational targeting restricts R-loop accumulation

**DOI:** 10.1101/2022.04.30.490097

**Authors:** Katarzyna B. Leszczynska, Monika Dzwigonska, Hala Estephan, Jutta Moehlenbrink, Elizabeth Bowler, Amato J. Giaccia, Jakub Mieczkowski, Bozena Kaminska, Ester M. Hammond

## Abstract

Local hypoxia occurs in most solid tumors and is associated with aggressive disease and therapy resistance. Widespread changes in gene expression play a critical role in the biological response to hypoxia. However, most research has focused on hypoxia-inducible genes as opposed to those which are decreased in hypoxia. We demonstrate that chromatin accessibility is decreased in hypoxia, predominantly at gene promoters and specific pathways are impacted including DNA repair, splicing and the R-loop interactome. One of the genes with decreased chromatin accessibility in hypoxia was *DDX5*, encoding the RNA helicase, DDX5, which showed reduced expression in various cancer cell lines in hypoxic conditions, tumor xenografts and in patient samples with hypoxic tumors. Most interestingly, we found that when DDX5 is rescued in hypoxia, replication stress and R-loop levels accumulate further, demonstrating that hypoxia-mediated repression of DDX5 restricts R-loop accumulation. Together these data support the hypothesis that a critical part of the biological response to hypoxia is the repression of multiple R-loop processing factors, however, as shown for DDX5, their role is specific and distinct.

## INTRODUCTION

Oxygen is an essential molecule in most biochemical reactions ^1^. During the lifetime of a cell, the oxygen concentration can fluctuate and therefore complex mechanisms have evolved to enable the cell to adapt and survive. Hypoxia is associated with pathological conditions such as cancer, stroke and cardiovascular disease and can lead to cellular damage ^2^. In cancer, regions of hypoxia arise when the poorly formed tumor vasculature fails to deliver sufficient oxygen to meet the high metabolic demand of rapidly proliferating cancer cells ^3^. The biological response to hypoxia involves a complex transcriptional program including the transactivation of hundreds of genes by the hypoxia inducible factors (HIFs) as well as the repression of specific processes, for example the DNA repair pathways ^4^. Hypoxia switches cellular metabolism from oxidative phosphorylation to aerobic glycolysis, yielding less energy. Therefore, wide-spread hypoxic-mediated repression of gene expression has been attributed to reserving cellular energy for vital processes required for cell survival. However, it is also plausible that in specific cases genes/pathways could be actively repressed as a critical part of the biological response to hypoxia.

R-loops are three-stranded nucleic acid structures consisting of an RNA/DNA hybrid and displaced single-stranded DNA, which originate during transcription ^5,6^. We recently demonstrated that R-loop levels increase in response to hypoxia (<0.1% O_2_) and that the RNA/DNA helicase, Senataxin (SETX), is induced in these conditions ^7^. An underlying mechanism for the hypoxia-mediated accumulation of R-loops has not been elucidated and the impact at specific gene loci has not been described. Interestingly, while the SETX helicase was induced in hypoxia, the expression of several other R-loop-associated factors (known as the R-loop interactome) was decreased, suggesting that the loss of these factors could contribute to the overall increase in R-loop levels ^7^. This finding led us to consider that an important biological response to hypoxia is to reduce the expression of R-loop related factors; however, the mechanism behind the global decrease of these factors in hypoxia remains unclear.

We and others have shown that hypoxia leads to widespread epigenetic changes, including a rapid increase in repressive histone methylation marks, for example H3K9me3, which is most likely attributed to impaired function of the oxygen-dependent histone lysine demethylases ^8–10^. Here, we investigated whether epigenetic changes affecting chromatin accessibility could contribute to the loss of the R-loop interactome in hypoxic conditions. As both the DNA repair and splicing pathways are repressed in hypoxic conditions and linked to R-loop formation and resolution, we also assessed if changes in chromatin accessibility might be responsible for the loss of these pathways ^4,11^. Using the assay for transposase-accessible chromatin with sequencing (ATAC-seq) we found a decrease in chromatin accessibility at the promoters of a number of genes encoding RNA processing factors, including R-loop-associated helicases. We focused on DDX5 which is an ATP-dependent DEAD/H-box RNA helicase, with diverse roles in mRNA splicing, co-activation of transcription factors, nonsense mediated decay and resolving R-loops ^12–19^. We show that restoring DDX5 levels in hypoxia results in further accumulation of replication stress and R-loops, suggesting that the repression of DDX5 in hypoxia restricts R-loop accumulation.

## RESULTS

To investigate the potential role of epigenetic changes in the regulation of the R-loop interactome in hypoxia we employed a murine glioblastoma model, GL261, which is known to develop hypoxic areas when grown orthotopically (**Fig. S1A**) ^20^. We assessed the changes in chromatin marks associated with a repressive heterochromatin state in hypoxia. GL261 cells were exposed to either moderate (1% O_2_) or severe (<0.1% O_2_) hypoxia and changes in chromatin marks were determined. As expected, we observed a marked increase in H3K9 and H3K27 trimethylation ^8,9,21^. However, we also observed an increase in H3K4 trimethylation, an activating histone modification. In all cases, increased methylation was most pronounced in severe hypoxia (<0.1% O_2_) and was reduced to near normoxic levels as quickly as 1 hour after reoxygenation, further emphasizing oxygen-dependent changes in chromatin (**Fig. 1A** and **Fig. S1B, C**). In addition, hypoxia (<0.1% O_2_) led to the loss of histone acetylation mark H3K27ac, which labels active promoters and enhancers (**Fig. S1B, C**).

**Fig. 1.**
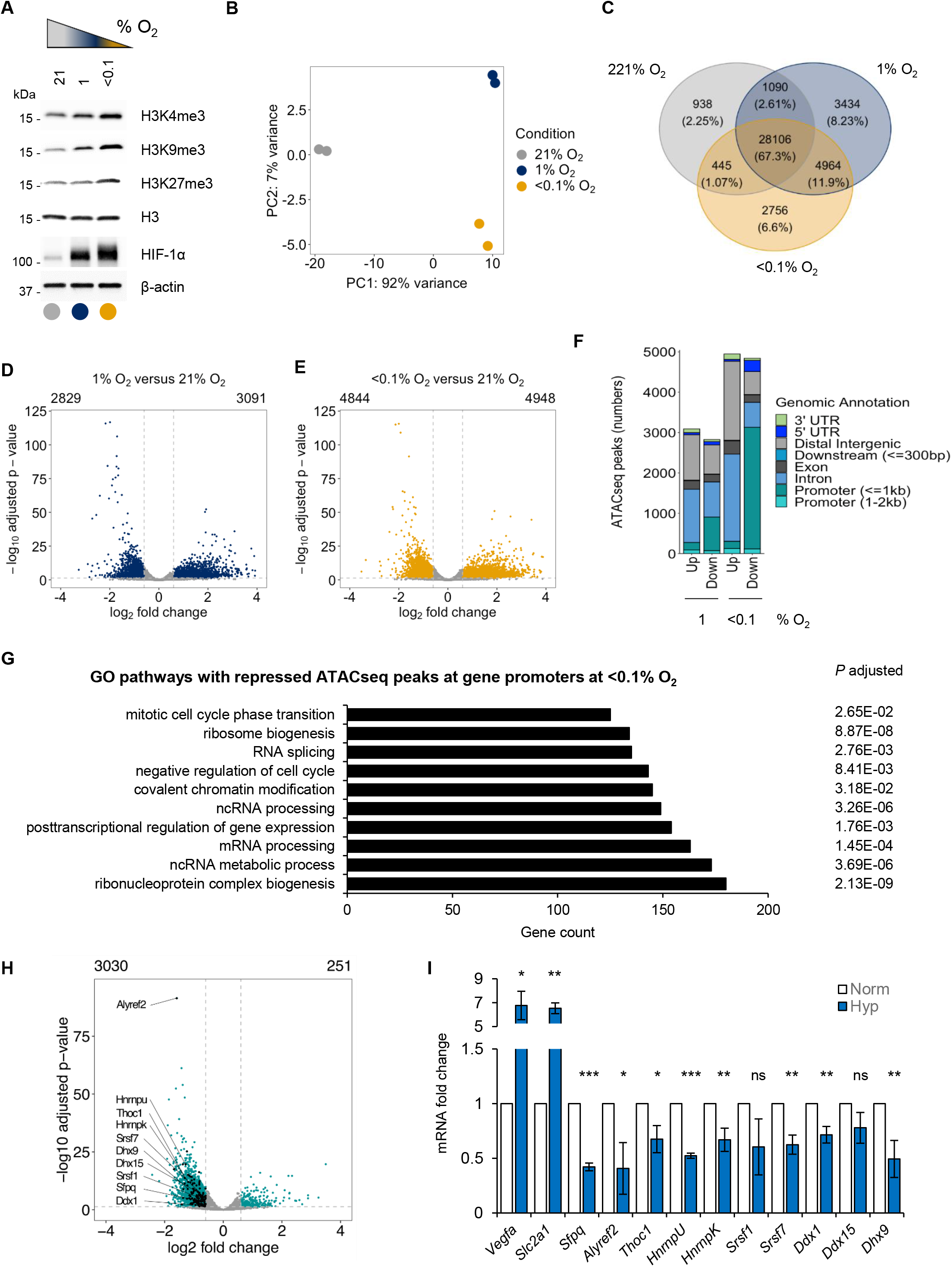
Oxygen-dependent chromatin alterations lead to a loss of promoter accessibility of numerous pathways, including RNA processing factors. **A**. GL261 cell line was exposed to 21, 1 or <0.1% O_2_ for 16 h and subjected to western blotting for the histone modifications indicated. HIF-1α was used as a hypoxia control. A representative western blot of three biological replicates is shown. **B**. GL261 cells were exposed to 16 h of normoxia (21% O_2_) or two conditions of hypoxia (1 and <0.1% O_2_). Samples were fixed and ATACseq carried out. Principal component analysis of ATACseq peaks for the three oxygen tensions is shown. **C**. Venn diagram showing hypoxia specific or common ATACseq peaks detected in all oxygen conditions from B. The percentage of peaks in the total amount of peaks is shown for each condition. **D**. A volcano plot showing differentially altered ATACseq peaks in 1% O_2_ versus 21% O_2_.Statistically significant peaks with FDR < 0.05 and |log2 fold change| ≥ 0.6 are marked in blue. **E.** A volcano plot showing differentially altered ATACseq peaks in <0.1% O_2_ versus 21% O_2_.Statistically significant peaks with FDR < 0.05 and |log2 fold change| ≥ 0.6 are marked in yellow. **F**. Differentially regulated ATACseq peaks from parts D and E annotated to distinct genomic regions at 1% or <0.1% O_2_ in relation to normoxic control. “Up” or “Down” marks significantly increased or decreased peaks, respectively, for each hypoxic condition in relation to normoxia. **G**. Top 10 Gene Ontology (GO) pathways (based on the highest gene count) associated with genes that had decreased ATACseq peaks at their promoters. Adjusted *p* value indicates statistical significance for each pathway shown. **H**. volcano plot showing differentially regulated ATACseq peaks at the gene promoters in cells treated with <0.1% O_2_, with significantly up-regulated 251 peaks and 3030 downregulated peaks. Statistically significant peak changes were annotated with green dots (FDR < 0.05 and |log2 fold change| ≥ 0.6). Black dots mark all significantly downregulated peaks for gene promoters of genes from the GO pathway “RNA splicing” (GO:0008380) and some names of these genes are indicated (full list in **Table S4**). **I**. qPCR validation of hypoxia-dependent repression of splicing factors indicated in H in GL261 cell line treated for 16 h at <0.1% O_2_. Mean ± standard deviation expression is shown from three biological repeats. *Vegfa* and *Slc2a1* were used as positive hypoxia-inducible controls. Statistical significance was calculated with two-tailed student t-test for each gene expression tested in hypoxia compared to normoxia (* *p*<0.05, ** *p*<0.01, *** *p*<0.001, ns = non-significant).

Since hypoxia-mediated increases in methylation marks could have a repressive or activating influence on gene expression, we investigated chromatin accessibility. We carried out ATACseq analysis on GL261 cells exposed to hypoxic (1 or <0.1% O_2_) or normoxic (21% O_2_) conditions. To prevent any reoxygenation-dependent changes in the chromatin that could occur during sample processing, we employed a modification of ATACseq that allows fixation of cells prior to lysis ^22^. Principal component analysis (PCA) showed a clear separation of ATACseq profiles between normoxic and the two hypoxic conditions (**Fig. 1B**). Overall, we detected 41,733 ATACseq peaks, of which 67.3% were detected in all conditions, 2.25% were specific to normoxia, 8.23% were specific to 1% O_2_ and 6.6% were specific to <0.1% O_2_ conditions (**Fig. 1C**). We then carried out a differential peak analysis between hypoxic and normoxic conditions and identified a total of 5,920 peaks with differential chromatin accessibility at 1% O_2_ and 9,792 peaks with differential chromatin accessibility at <0.1% O_2_, in relation to the normoxic control (**Fig. 1D, E**). Of these, 3,091 peaks were significantly increased, and 2,829 peaks were significantly decreased at 1% O_2_, while 4,948 peaks were significantly increased, and 4,844 peaks were significantly decreased at <0.1% O_2_ (**Fig. 1D, E** and **Table S1**). Overall, severe hypoxia led to a greater number of significant ATACseq peak changes, which reflects the oxygen-dependent changes in histone methylation that are known to contribute to the chromatin accessibility (**Fig. 1A** and **Fig. S1B, C**) ^23,24^. Next, we carried out genomic annotation of the differentially regulated ATACseq peaks in hypoxia (1 and <0.1% O_2_) (**Fig. 1F** and **Fig. S1D**). One of the most notable differences between the response to moderate (1% O_2_) and severe (<0.1% O_2_) hypoxia was the number of ATACseq peaks lost at promoter regions, with nearly 1000 peaks lost in 1% O_2_ and over 3000 peaks lost in <0.1% O_2_. The oxygen dependency was further emphasized when a ratio between a percentage of decreased peaks at specific genomic annotations in severe versus moderate hypoxia was evaluated (**Fig. S1E, F**), where again the highest percentage of decreased ATACseq peaks at promoter regions was in response to severe hypoxia compared to moderate hypoxia (**Fig. S1F** and **Table S2**). Moreover, the majority of repressed promoter peaks identified in moderate hypoxia (789 out of 852) were included in the peaks repressed in severe hypoxia (**Fig. S1G**). However, severe hypoxia (<0.1% O_2_) repressed additional promoters in comparison to moderate hypoxia (1% O_2_), demonstrating the oxygen dependency of this process. Since severe hypoxia induced the greatest loss of accessible promoters, we performed a functional enrichment analysis of the genes with significantly decreased ATACseq peaks at promoter regions in severe hypoxia using three databases: Gene Ontology (GO), Kyoto Encyclopedia of Genes and Genomes (KEGG) and Reactome ^25–27^. We identified several signaling pathways associated with the peaks decreased in severe hypoxia, many of which were related to RNA metabolism and processing, including ribonucleotide complex biogenesis, ncRNA metabolic process or mRNA processing and splicing. The top ten significantly downregulated pathways from GO analysis based on the highest gene count associated with each pathway are shown (**Fig. 1G**, full list of significantly associated pathways **Table S3**). In order to ensure robust ATACseq results, we employed two additional data normalization approaches, as it was recently shown that choice of analysis can impact differential chromatin accessibility results ^28^. Using two alternative normalization pipelines, we found that the impact of hypoxia on the chromatin accessibility remained qualitatively similar as determined with the initial DEseq2 analysis. We observed changes in the number of significantly up-regulated or down-regulated ATACseq peaks between every method used; however, the effect on the repression of accessible promoters in conditions of severe hypoxia, including genes affecting RNA processing and splicing was observed with all methods (**Fig. S2**).

To validate the results of our ATACseq analysis, we focused on genes involved in splicing, as components of this pathway have previously been shown to be transcriptionally repressed in hypoxia ^11^. From the list of the spliceosome genes, we identified a number of splicing factors as having decreased ATACseq peaks in hypoxia (**Table S4**) and confirmed that 10 of them were repressed at the mRNA level (**Fig. 1H, I**). This was in contrast to the mRNA of known HIF targets, *Slc2a1* or *Vegfa*, which are induced in response to hypoxia ^29^. Moreover, the functional analyses of decreased ATACseq peaks in gene promoter regions led to the identification of other pathways previously reported as decreased in hypoxia, including the DNA repair pathways; base excision repair, homology-directed repair, double strand break repair and non-homologous end-joining (Reactome analysis, **Table S3**) ^4^. Overall, our data identifies a global mechanism of hypoxia-induced repression of multiple pathways linking previously reported mRNA repression with decreased chromatin accessibility.

Loss of chromatin accessibility at the promoters of genes coding for RNA-processing factors supports our hypothesis that the repression of genes involved in R-loops could be mediated by epigenetic changes, as many of these genes have been identified in the R-loop interactome. The RNA and RNA/DNA helicases *Ddx1, Ddx3x, Ddx5, Ddx18, Ddx21, Ddx27, Ddx39b, Dhx9, Dhx15, Srsf1, Xrn2* and *Prkdc* are all components of the R-loop interactome and were all found to have reduced chromatin accessibility in their promoters in hypoxia (**Fig. S3A** and **Table S5**) ^30^. Of these helicases, we have shown previously that *DHX9* mRNA expression is repressed in hypoxia (<0.1% O_2_) therefore supporting a link between these ATACseq data and gene expression in hypoxia (**Fig. S3B**) ^7^. In contrast, chromatin accessibility did not significantly change at the *SETX* gene promoter (**Fig. S3C**), consistent with the recently identified increase in *SETX* expression in response to hypoxia (<0.1% O_2_) ^7^. We then asked whether the chromatin accessibility changes in hypoxia detected with ATACseq were mirrored by expression of identified R-loop and spliceosome interacting genes. We investigated this hypothesis using the transcriptional atlas of human glioblastoma, where RNAseq analysis was performed on samples micro-dissected from specific regions within the glioblastoma, including well oxygenated “Microvascular Proliferation” regions, as well as “Pseudopalisading Cells Around Necrosis”, which are likely to be hypoxic (Ivy Glioblastoma Atlas Project; http://glioblastoma.alleninstitute.org/) ^31^. A heatmap generated from the z-scores of this data showed a high expression of known hypoxia-inducible genes (*VEGFA, SLC2A1, CA9, LDHA, BNIP3, BNIP3L* and *LDHA*) in Pseudopalisading Cells Around Necrosis. In contrast, the expression of numerous genes from the GO Splicing pathway (GO:0008380) as well as the R-loop interactome, whose promoters had decreased ATACseq signal in hypoxic conditions in our dataset, were downregulated at the mRNA level in human glioblastoma samples, emphasizing the clinical relevance of our ATACseq findings (**Fig. 2A**). One of the novel hypoxia-repressed targets identified here in ATACseq analysis and confirmed with the IvyGAP dataset was *DDX5*, a RNA helicase that was recently reported as a crucial factor in resolving R-loops (**Fig. 2C** and **Fig. S3D**) ^15–19^.

**Fig. 2.**
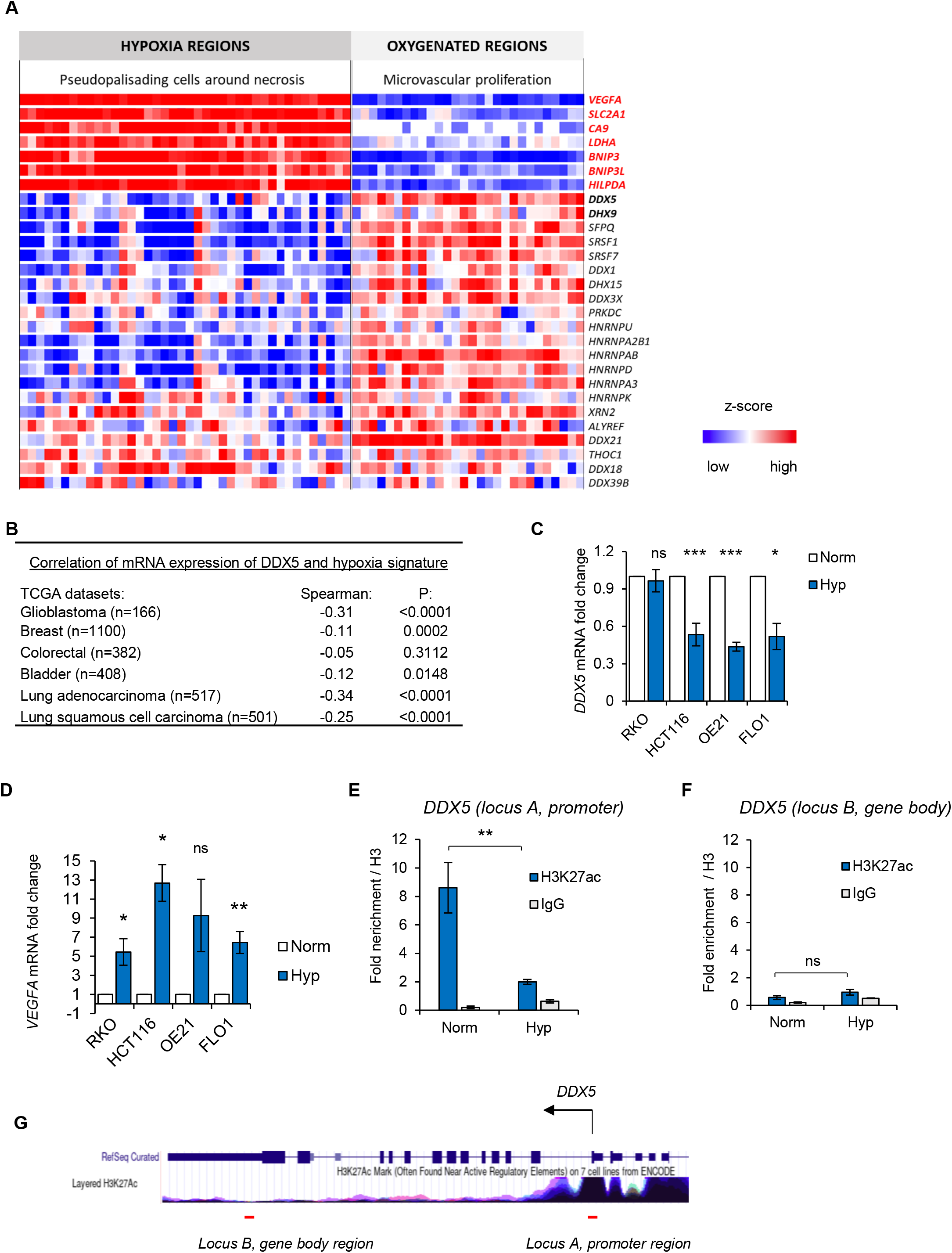
Loss of chromatin accessibility in hypoxia is mirrored by gene expression changes, including the loss of mRNA expression of *DDX5*. **A**. A heatmap showing a z-score expression of hypoxia markers as well as RNA processing factors in two anatomical regions of glioblastoma, including “microvascular proliferation” region and “pseudopalisading cells around necrosis”. Data was accessed via IvyGAP Glioblastoma project at http://glioblastoma.alleninstitute.org/ and selected genes are shown. Each column represents a separate sample for a given region and rows show expression of particular genes in these samples. **B**. Expression of *DDX5* (mRNA) was correlated with the hypoxia metagene signature in the indicated TCGA cancer patient cohorts: breast invasive carcinoma, colorectal adenocarcinoma, glioblastoma, bladder urothelial carcinoma, lung adenocarcinoma and lung squamous cell carcinoma. The numbers of patient samples are shown in brackets. Spearman’s rank correlation coefficient and *p* values are shown for the Log10 median expression of *DDX5* and hypoxic signature. **C, D**. The indicated cell lines were exposed to hypoxia (<0.1% O_2_) for 18 h. Expression of *DDX5* (C) and *VEGFA* (D) was determined by qPCR and normalized to *18S*. Statistical significance was calculated with two-tailed student t-test for each cell line based on three biological replicates (* *p*<0.05, ** *p*<0.01, *** *p*<0.001, ns = non-significant). **E, F**. HCT116 cells were exposed to 16 h of hypoxia (<0.1% O_2_) followed by chromatin immunoprecipitation with H3, H3K27ac and IgG antibodies and qPCR analysis. Fold enrichment for H3K27ac and IgG normalized to total H3 enrichment is shown as mean ± standard deviation from three independent experiments. Statistical significance was calculated with two-tailed student t-test (** *p*<0.01, ns = non-significant). **G**. The *DDX5* locus taken from the UCSC genome browser (https://genome.ucsc.edu/) at the human GRCh37/hg19 genome assembly is shown with a track underneath showing the layered H3K27ac profile across the *DDX5* gene. Red lines underneath the track indicate the binding site of ChIP-qPCR primers amplifying the promoter region (locus A) and gene body region (locus B) used in ChIP-qPCR analysis in E and F.

To explore potential links between DDX5 and hypoxia in cancer, we asked if *DDX5* expression and a validated hypoxia signature correlated in TCGA cancer patient cohorts (glioblastoma, breast, colorectal, bladder, lung adenocarcinoma and lung squamous cell carcinoma). We found a significant inverse correlation between *DDX5* mRNA and the hypoxic signature in five out of the six tested datasets (**Fig. 2B** and **Fig. S4**). These findings suggest that DDX5 mRNA expression is repressed in hypoxic tumors. This association was further verified in cancer cell lines using qPCR. In three of the tested cell lines (HCT116, OE21, FLO1), we observed significant repression of *DDX5* mRNA (approximately 50% of the normoxic control) in contrast to the HIF-1 target *VEGFA* which was induced in hypoxia (**Fig. 2C, D**). To test whether *DDX5* mRNA repression could be directly linked with chromatin accessibility changes at the *DDX5* promoter, we carried out ChIP-qPCR for H3K27ac. H3K27ac, which labels active gene promoters and enhancers, was significantly enriched at the promoter region of *DDX5* in normoxic conditions. However, this enrichment was lost in response to hypoxia (**Fig. 2E, G**). In contrast, H3K27ac was not enriched in the *DDX5* gene body distal to the promoter, in normoxic or hypoxic conditions (**Fig. 2F, G**). Overall, these data support our ATACseq suggesting that *DDX5* expression might be repressed at the transcriptional level, via restriction of the accessible chromatin at the promoter of *DDX5* gene and loss of active chromatin marks, including H3K27ac.

Next, we asked if changes in chromatin accessibility and mRNA expression of *DDX5* correlated with changes in protein expression. DDX5 protein was significantly decreased in cells exposed to hypoxia (<0.1% O_2_) and was quickly reversed when oxygen was restored to normoxic levels reflecting the changes to the histone modification marks during hypoxia and reoxygenation (**Fig. 3A** and **Fig. S1C**). DHX9 was also included and found to decrease in response to hypoxia (**Fig. 3A)**. As the HIF-1 transcription factor is known to regulate chromatin in hypoxic conditions, we further investigated the oxygen-dependency of DDX5 repression and possible HIF involvement, using a range of hypoxic conditions (<0.1 or 2% O_2_) as well as the hypoxia mimetics CoCl_2_ and DFO ^32^. Exposure to 2% O_2_, CoCl_2_ or DFO, had little or no effect on DDX5 expression, indicating that the repression of DDX5 is unlikely to be dependent on HIF-mediated signaling (**Fig. S5A, B)**. To confirm this hypothesis, RKO cells lacking HIF-1α were exposed to hypoxia and the levels of DDX5 determined compared to the control cell line. As expected, DDX5 levels decreased in both cell lines demonstrating that this occurs independently of HIF-1α stabilization (**Fig. 3B, C)**. As DDX5 expression has not previously been linked to the response to hypoxia, we investigated its expression in a panel of human cell lines. In all the tested cancer cell lines, including lung, esophageal, colorectal, bladder, bone, and glioma cells, DDX5 protein was repressed in response to hypoxia (<0.1% O_2_) (**Fig. 3D-I** and **Fig. S5C-E**). Notably, a similar change occurred in non-cancer MRC-5 cells, indicating that the hypoxia-mediated repression of DDX5 is not restricted to cancer cells (**Fig. S5F**).

**Fig. 3.**
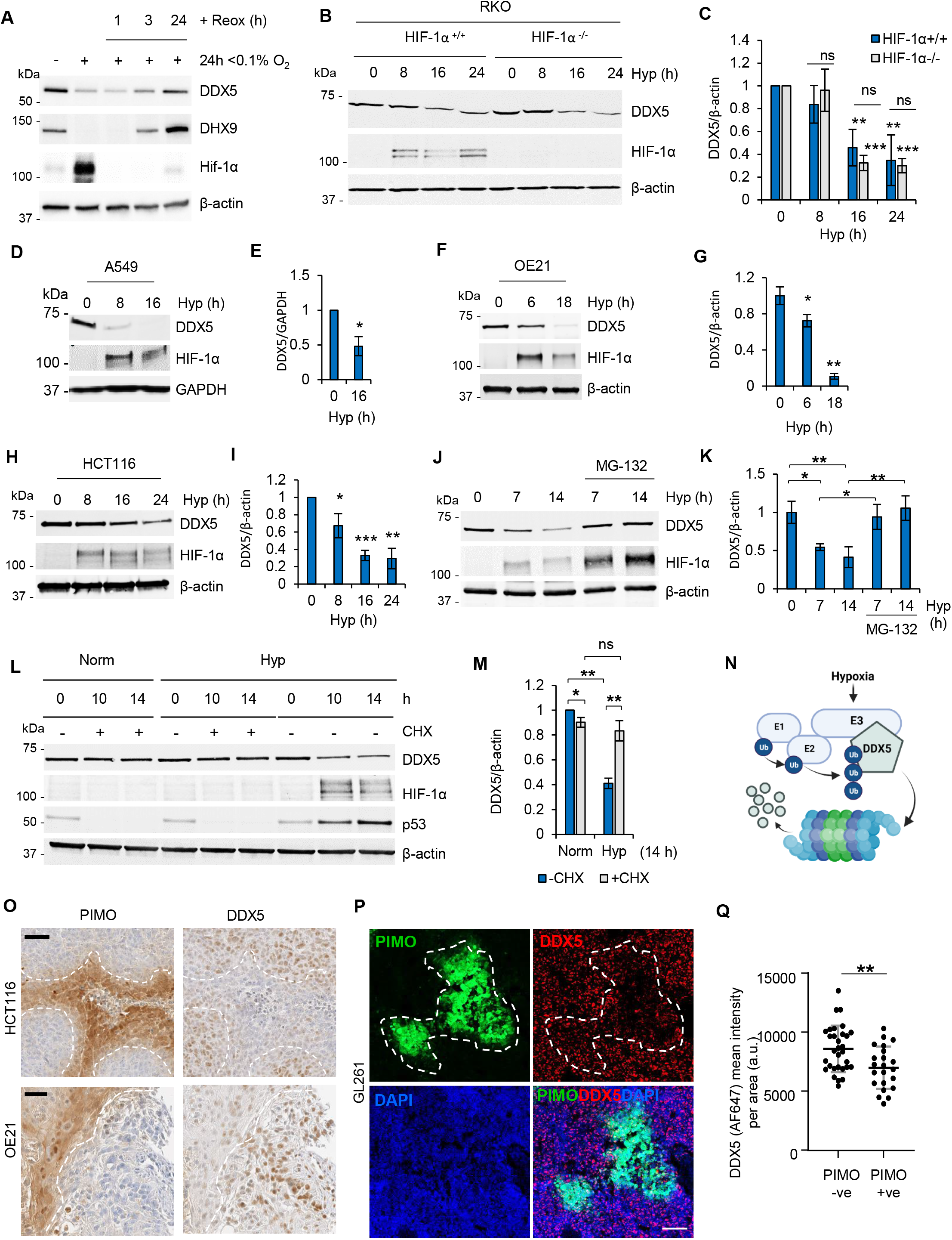
Hypoxia leads to the active repression of DDX5 protein. **A**. GL261 cells were exposed to hypoxia (Hyp, 0.1% O_2_) for 24 h followed by reoxygenation (to 21% O_2_) for the indicated times following hypoxia. Western blotting was carried out for the antibodies shown (representative of three independent experiments). **B, C**. RKO^HIF-1α+/+^ and RKO^HIF-1α-/-^ were exposed to hypoxia (<0.1% O_2_) for the times indicated and subjected to western blotting with the antibodies shown (representative of at least three biological replicates). Densitometry is shown in part C. **D-I**. A549, OE21 and HCT116 cell lines were exposed to hypoxia (Hyp, 0.1% O_2_) for the times indicated followed by western blotting for DDX5 as well as HIF-1α (hypoxic marker) and β-actin (loading control). Representative blots from at least three independent experiments are shown. Densitometry for the biological replicates showing hypoxia-dependent repression of DDX5 is shown. **J, K**. HCT116 cells were exposed to hypoxia (Hyp, <0.1% O_2_) and MG-132 (5 µM) as indicated and western blotting was carried out with the antibodies shown. Densitometry for three biological replicates is shown in part K. **L, M**. HCT116 cells were treated with 25 µg/ml cycloheximide (CHX) in hypoxic (Hyp, <0.1% O_2_) or normoxic (Norm, 21% O_2_) conditions for the times indicated and analyzed with western blotting. Densitometry for three biological replicates is shown in part M. **N.** A scheme showing hypothesis of hypoxia-inducible degradation of DDX5 via proteasome. The scheme was drawn with BioRender.com. **O.** Sequential sections from HCT116 and OE21 tumor xenografts using the cell lines indicated (HCT116 or OE21) were stained for hypoxia with anti-pimonidazole hyoxyprobe-1 (PIMO) and DDX5 antibodies. Nuclei were counterstained with hematoxylin Scale bar = 50 μm. The outline of PIMO-positive (brown) staining is shown by the dashed white line. **P.** PFA-fixed GL261 tumor xenografts were sectioned and subjected to immunofluorescent staining for hypoxia (PIMO) and DDX5. Nuclei were counterstained with DAPI. Scale bar = 100 μm. The outline of PIMO-positive (FITC) staining is shown by the dashed white line. **Q.** Mean fluorescence intensity ± standard deviation of DDX5 signal (labeled with Alexa Fluor-647) in GL261 tumors from part **P** was measured in hypoxic (PIMO+ve) and normoxic (PIMO-ve) image areas using ZEN2 software (Zeiss). A total of 22 hypoxic areas and 31 normoxic areas were analyzed and two-tailed non-parametric Mann-Whitney test shows statistical significance (*p* value 0.007, **)

DDX5 is known to interact with its paralog DDX17 and depletion of DDX5 has been shown to upregulate DDX17 ^14^. Therefore, we asked whether loss of DDX5 expression in hypoxia would lead to increased expression of DDX17. When we depleted *DDX5* using siRNA, the levels of DDX17 increased significantly in normoxic conditions confirming the compensatory relationship between DDX17 and DDX5 (**Fig. S5G, H**). However, in hypoxic conditions, where DDX5 is repressed, the expression of DDX17 did not change, indicating that this compensatory relationship does not occur when DDX5 is repressed by hypoxia. Together, these data suggest that DDX5 is repressed in an oxygen-dependent manner and this event does not result in increased expression of its paralog, DDX17.

Next, we tested the involvement of the proteasome pathway in the downregulation of DDX5 and confirmed that DDX5 protein was degraded via the proteasome in hypoxia by co-treating cells with a proteasome inhibitor, MG-132 (**Fig. 3J, K**). We then went on to investigate the half-life of the DDX5 protein in hypoxia and in normoxia and surprisingly, found that blocking global protein translation with cycloheximide rescued DDX5 expression in hypoxia (**Fig. 3L, M**). Using emetine, as an alternative agent to block translation, we confirmed that hypoxia-mediated downregulation of DDX5 could be reversed by blocking translation (**Fig. S5I, J**). These data suggest that active translation is required in hypoxic conditions to decrease the levels of DDX5 and led to the hypothesis that an E3 ligase induced in hypoxia could target DDX5 for proteasomal degradation (**Fig. 3N**). In support of this hypothesis, we observed a partial, but consistent, rescue of DDX5 repression in hypoxia in the presence of a neddylation inhibitor, MLN4924, indicating the potential involvement of an E3 ubiquitin ligase/s, from the Cullin-RING type family (**Fig. S5K**).

Finally, we analyzed tumor xenografts (HCT116, OE-21 and GL261) for both hypoxia (using pimonidazole) and DDX5 expression. In all three tumor types, an inverse correlation was observed between hypoxia (pimonidazole positive) and DDX5 expression, confirming that hypoxia represses DDX5 protein expression *in vivo* (**Fig. 3O-Q**). Together, these data suggest that expression of DDX5 is regulated at multiple levels, including the decreased accessibility of the chromatin at the promoter of *DDX5*, decreased *DDX5* mRNA and enhanced protein degradation in hypoxia.

We hypothesized that the widespread repression of R-loop related factors in hypoxia could in part explain the increase in R-loops in hypoxic conditions ^7^. We investigated whether rescuing DDX5 in hypoxia could reduce the accumulation of R-loops in hypoxia with the obvious caveat that rescuing the expression of a single component of the R-loop interactome could have little impact. However, as R-loops are transcription dependent and DDX5 is a transcription co-factor, we first determined if rescuing DDX5 in hypoxia increased this activity. To begin, we measured changes in global transcription using 5’EU incorporation in cells exposed to hypoxia (<0.1% O_2_) with and without DDX5 over-expression. Rescuing DDX5 expression in hypoxia in both A549 and HCT116 cells had no impact on 5’EU incorporation (**Fig. 4A, B** and **Fig. S5A-C**). Furthermore, as DDX5 is a known co-factor of E2F1 and we have previously shown that some E2F target genes are repressed in hypoxia ^33^, we also overexpressed DDX5 in hypoxia and tested expression of known E2F target genes: MCM2, MCM4, MCM5, MCM6, MCM7 and CDC6. As expected, we observed a clear decrease in protein expression of these E2F target genes in hypoxia, however levels were not rescued when DDX5 was restored in hypoxia (**Fig. S5D)**. We then investigated the effect of restoring DDX5 expression in hypoxia on R-loop accumulation. DDX5 was expressed in cells and exposed to hypoxia (<0.1% O_2_), and used the catalytic-dead mutant RNase H1^D210N^ to visualize R-loops ^34,35^. In addition, we included a DDX5 helicase mutant (DDX5^NEAD^) and DHX9 as controls. Surprisingly, we found that rescuing DDX5 in hypoxia led to a significant accumulation of R-loops and importantly that this was dependent on DDX5 helicase activity (**Fig. 4C-E**). In contrast, when we restored the expression of DHX9, R-loop levels were significantly reduced both in normoxia and in hypoxia. To confirm this surprising finding, we verified the role of DDX5 in R-loop accumulation in hypoxia in a second cell line, HCT116. Again, we found that when DDX5 expression was restored in cells exposed to hypoxia, the levels of R-loops increased further (**Fig. S5E-G**). These data demonstrate that in hypoxic conditions, the restoration of DDX5-mediated helicase activity increases R-loop levels and that this is specific to the DDX5 helicase, as over-expression of DHX9 abrogated R-loops. Increasing R-loops in hypoxia would be predicted to lead to a further accumulation of replication stress ^7^. To investigate this hypothesis, we determined the impact of rescuing DDX5 expression in hypoxia on the levels of RPA foci as a marker of replication stress. A significant increase in the number of RPA foci per cell was seen in hypoxic cells with over-expressed DDX5 (**Fig. 4F, G**). Overall, these data suggest that hypoxia-mediated loss of DDX5 expression acts to restrict replication stress through decreased R-loop accumulation.

**Fig. 4.**
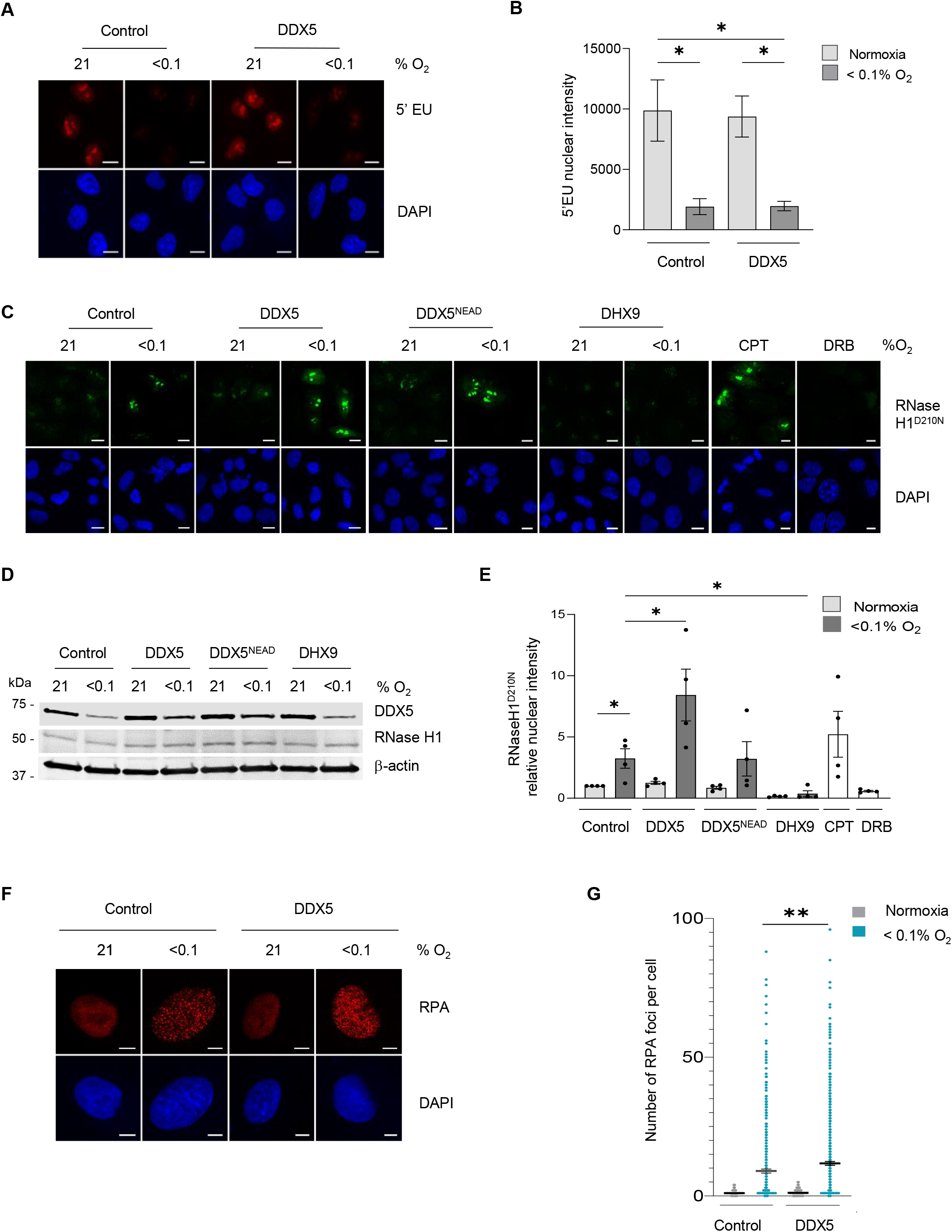
Rescuing DDX5 expression in hypoxia leads to accumulation of R-loops. **A.** A549 cells were transfected with myc-tagged DDX5 or control vector and exposed to hypoxia (8 h) with 5’EU (0.5 mM) added for the final hour. Staining for 5’EU was then carried out. Representative images are shown (scale bar = 10 μm). 5’EU staining in red, DAPI (blue) shows the nucleus. **B.** Nuclear Intensity of 5’EU staining from A was determined. Data represents the mean expression and SEM from three independent experiments. Statistical significance was calculated with an unpaired student t-test for each indicated condition (* *p*<0.05). **C.** A549 cells were co-transfected with RNase H1^D210N^ together with myc-tagged DDX5, myc-tagged DDX5^NEAD^, or with DHX9 and exposed to hypoxia for 18 hours. CPT (10 μM, 20 min) or DRB (100 μM, 1 h) were used as a positive and negative controls for R-loops, respectively. Representative images for R-loop visualization with V5-RNase H1^D210N^ fluorescence are shown (scale bar = 10 μm). V5 fluorescence is shown in green, DAPI (blue) shows the nucleus. **D.** Western blot analysis confirms expression of DDX5 and RNase H1^D210N^ across conditions shown in part C. **E.** Quantification of fluorescent intensity from part C. A minimum of 100 cells were included per condition. Data represent the mean expression and SEM from four independent experiments. Statistical significance was determined with an unpaired student t-test for each indicated condition (* *p*<0.05). **F.** A549 cells were transfected as indicated and staining for RPA foci was then carried out. Representative images are shown (scale bar = 10 μm). RPA staining in red, DAPI (blue) shows the nucleus. **G.** Quantification of the number of foci per cell in part F was determined. Data represents the mean expression and SEM from three independent experiments, each dot represents a cell and a minimum of 100 cells were imaged per treatment. Statistical significance was calculated with an unpaired student t-test for each indicated condition (** *p*<0.01).

## DISCUSSION

We demonstrate that many genes involved in R-loop formation and resolution (the R-loop interactome) are repressed in hypoxic conditions, with the notable exception of SETX. Our ATACseq analysis suggests that in most cases the repression of the R-loop interactome correlates with decreased chromatin accessibility, likely leading to reduced transcription. Changes in chromatin accessibility in hypoxia have been previously characterized in hypoxia-induced cardiac damage, adaptation to high-altitude, or cancer cell lines, and both chromatin condensation and increased open chromatin at various loci, including hypoxia-responsive genes, were reported ^32,36–38^. While changes to chromatin in hypoxia have been largely attributed to changes in chromatin modifying enzymes, it is also likely that changes in transcription factor activity in these conditions may impact chromatin and ultimately accessibility/gene expression ^39^. Recently, ATACseq was also used to demonstrate that the type I interferon pathway is downregulated in hypoxia, and that this is dependent on decreased chromatin accessibility at genes with motifs for STAT1 and IRF3 ^40^. Here, we found further pathways downregulated through this mechanism in hypoxia, including DNA repair, splicing and the R-loop interactome. It is plausible that the reduced expression of genes in all of these pathways contributes to R-loop levels in hypoxia.

Evidence to support the hypothesis that the repression of these genes is part of the biological response to hypoxia as opposed to an energy saving measure comes from our finding that in the case of DDX5, multiple mechanisms exist in hypoxia to reduce expression. Our data suggests increased protein degradation in hypoxia via a hypoxia-inducible E3 ubiquitin ligase, potentially as a means of quickly reducing DDX5 levels in addition to the loss of chromatin accessibility. In addition, the induction of *miR-462* and *-731* in hypoxia in a HIF-dependent manner has been shown to repress DDX5 in zebrafish, further supporting our finding that multiple mechanisms exist to reduce DDX5 in hypoxia ^41^. Surprisingly, whilst we predicted that loss of the R-loop interactome contributed to R-loop levels in hypoxia, we found that this was not a general response as rescuing DDX5 increased R-loops whilst the opposite effect was detected when DHX9 was restored. Notably our studies have focused on the global levels of R-loops and do not address changes at specific genomic locations.

Why restoring DDX5 expression in hypoxia leads to further R-loop accumulation above the level observed in hypoxia is unclear. Previous reports have all shown that expression of DDX5 leads to R-loop resolution i.e., a decrease in R-loop levels ^15–19^. To our knowledge, expression of DDX5 in hypoxia is the first example of a context where DDX5 has been linked to R-loop accumulation. One hypothesis is that this is related to the changes in splicing also observed in these conditions, as recruitment of the spliceosome onto nascent mRNA attenuates R-loop formation ^42^. In support of this, over-expression of DHX9 in the absence of key splicing factors has been shown to increase R-loop accumulation ^43^. However, when we restored DHX9 in hypoxia, we found an almost complete abrogation of R-loops, although notably this also occurred in the normoxic samples, making the impact of DHX9 in hypoxia difficult to interpret.

A caveat of having chosen DDX5 as an exemplar for the R-loop interactome is that DDX5 has numerous functions which are independent of R-loops ^12–14,44–46^. For example, it is plausible that repression of DDX5 could be a part of the biological adaption to reduced oxygen levels as shRNA-mediated depletion of DDX5 in lung cancer cells leads to reduced oxygen consumption through changes in mitochondrial respiration, oxidative phosphorylation and reduced intracellular succinate ^47^. Therefore, reduction of DDX5 expression in hypoxia may alter numerous alternative pathways, which in turn may also indirectly affect R-loop levels, making the direct effect of DDX5 on R-loop levels difficult to determine.

One of the aspects of DDX5 which intrigued us was the finding that is overexpressed or amplified in a wide range of cancers ^48^. Initially our finding that hypoxia leads to loss of DDX5 seemed contradictory to the finding that it is over-expressed in cancers. However, when considered together, this demonstrates that whilst hypoxic tumors express less DDX5, the levels are increased compared to normal tissues. This in turn suggests that the *DDX5* gene amplification in cancers could occur to compensate for the repressive effect of hypoxia on DDX5. Together our data highlight the significant energy-consuming processes a hypoxic cell employs to turn off the expression of specific genes such as DDX5 and, not surprisingly, that cancers circumvent this biological response.

## METHODS

### Cell lines and reagents

HCT116, RKO, U87, A549, MCF-7, MDA-231, DLD1, H1299, WI-38 and MRC5 cells (from ATCC) were cultured in DMEM with 10% FBS. OE21 and FLO1 cell lines were obtained from PHE culture collections and cultured in RPMI and DMEM, respectively, both supplemented with 10% FBS. RKO^HIF-1α-/-^ cells were obtained from Dr Denise Chan, UCSF. GL261 mouse glioma cells stably expressing either pEGFP-N1 or tdTomato (luc+tdT+) were cultured in DMEM with 10% FBS and antibiotics (50 U/ml penicillin, 50 mg/ml streptomycin) ^49,50^. NBE1 cells were cultured in DMEM F12 advanced, glutamax and 1% FBS while ARPE-19 cells were cultured in DMEM F12 medium supplemented with 10% FBS (cells from Prof. Geoff Higgins, University of Oxford). ReNCell CX, ReNCell VM (from Prof. Eric O’Neill, University of Oxford). HCT116 p53+/+ and p53-/- cell lines were a gift from Prof. Vogelstein (Johns Hopkins University, Maryland, USA). DharmaFECT 1 transfection reagent (Dharmacon) was used for siRNA transfections according to the manufacturer’s instructions; sequences are available in the SI. pSG5-myc, pSG5-mycDDX5 and pSG5-mycDDX5-NEAD plasmids were a kind gift from Prof. Frances Fuller-Pace (University of Dundee) and DHX9 plasmid from Dr Natalia Gromak (University of Oxford). ppyCAG_RNaseH1_D210N_V5 plasmid was obtained from Addgene (#111904). Plasmid transfection was carried out using jetPRIME® (Polyplus) transfection reagents according to the manufacturer’s protocol.

### Hypoxic conditions and drug treatment

Cells were incubated at <0.1% O_2_ in a Bactron chamber (Shel Lab) or at <0.1%-2% O_2_ in a M35 Hypoxystation (Don Whitley) hypoxic chamber in the presence of 5% CO_2_. The following drugs were used: 25 µg/ml Cycloheximide, 20 µM Emetine, 5 µM MG-132, 2 µM MLN-4924, 1 µM Ruxolitinib, 100 µg/ml RNAseA, 1×10^3^ U/ml IFNα, 100 µg/ml INFγ, 100 µM 5,6-dichloro-1-β-d-ribofuranosylbenzimidazole (DRB) and 10 µM Camptothecin (CPT)

### Western blotting

Cells were washed in PBS and lysed in SDS lysis buffer (10 mM Tris-Cl, pH 7.5, 0.1 mM EDTA, 0.1 mM EGTA, 0.5% SDS, 0.1 mM β-mercaptoethanol, protease/phosphatase inhibitors). After blocking in LiCOR blocking buffer the following primary antibodies were used: DDX5 (Pab204, gift from Prof. Frances Fuller-Pace; clone204, Sigma-Aldrich), DHX9 (ab26271, Abcam), DDX17 (SQQ-K14, gift from Prof. Frances Fuller-Pace, Dundee, UK), p53 (DO-I, Santa Cruz), H3K4me3 (07-473, Millipore), H3K9me3 (ab8898, Abcam), H3K27me3 (07-449, Millipore), H3 total (ab1791, Abcam) β-actin (AC-15, Santa Cruz), HIF-1α (610958, BD Biosciences), and GAPDH (6C5, Novus Biologicals). IRDye® 680 or IRDye® 800 secondary antibodies were used and the Odyssey infrared system (LI-COR) to visualize western blots.

### Chromatin immunoprecipitation (ChIP)

ChIP method was adapted from Cook *et al*., with some modifications ^51^. 2×10^6^ HCT116 cells were used per each immunoprecipitation (IP) reaction and 15 units of micrococcal nuclease (MNase, Worthington) was used to digest the chromatin for the ChIP reaction. 4 ug of antibody was used for each IP reaction: H3 (ab1791, Abcam), H3K27ac (#8173) and normal rabbit IgG (#2729S, Cell Signaling Technology) together with 20 μl of pre-washed Protein A/G Dynabeads Chromatin digestion was assessed using TapeStation electrophoresis system (Agilent Technologies). Desired DNA digestion yielded a mono- and poly-nucleosome pattern with a range of DNA fragments 100-1000 bp. Purified input and ChIP samples were subjected to qPCR with Applied Biosystems™ Fast SYBR™ Green Master Mix (4385612, Applied Biosystem) according to the manufacturer’s instructions using QuantStudio 12 K Flex Real-Time PCR System. Primers were used for DDX5 promoter locus (forward primer: ATGTTCCTTCGTCTGCCTCGA, reverse primer: CTTGCTTTTTGTGTGGGGATT) and DDX5 gene body locus (forward primer: TGAAAACACTGCCTGCATTTT, reverse primer: AATTGCAGAAATGACTGCAGT). Percent of input was determined for total H3, H3K27ac and IgG control IP samples (2^-(sample Ct– adjusted input Ct)). Fold enrichment for H3K27ac and IgG was determined relative to H3. Mean and standard deviation from three biological replicates was plotted.

### 5’EU staining

Cells were incubated with 5’ethynyluridine (EU) (0.5 mM) then fixed in 4% paraformaldehyde for 10 min and permeabilized in 0.1% Triton-X 100 for 10 min. Click-iT Alexa Fluor 647 labeling kit (ThermoFisher) was used to measure nascent transcription in the entire nucleus. Images were captured using a LSM710 confocal microscope (Carl Zeiss, Germany). Image processing and analysis were performed using Image J software. The 5’EU mean nuclear intensity signal was determined using the Image J plugin/algorithm, measured as previously ^7^. A minimum of 100 cells were included per condition.

### RPA foci staining

Cells transfected with DDX5^WT^ were fixed in 4% paraformaldehyde for 10 min, permeabilized in 0.1% Triton-X 100 for 10 min and blocked with 2% BSA in 0.1% Tween-20 in PBS for 1 hour. Cells were then incubated with RPA32 antibody (2208, Cell Signaling) for 2 hours at 37°C followed by Alexa fluor rat 594 secondary antibody (A-11007, Invitrogen) for 1 hour at room temperature. Coverslips were mounted using mounting medium Prolong® Gold with DAPI (P36962, Invitrogen). Images were captured using a LSM710 confocal microscope (Carl Zeiss, Germany). Image processing and analysis were performed using Image J software. R-loop quantification Cells transfected with RNase H1^D21ON^ V5 were extracted in ice-cold PBS/0.2% Triton-X 100 for 2 min, fixed in 4% paraformaldehyde for 10 min, permeabilized in 0.1% Triton-X 100 for 10 min and blocked with 1% BSA /FBS in PBS for 1 hour. Cells were then incubated with V5 antibody (R96025, ThermoFisher Scientific) followed by Alexa fluor mouse 488 secondary antibody (A11029, Invitrogen) for 1 hour at room temperature. Coverslips were mounted using mounting medium Prolong® Gold with DAPI (P36962, Invitrogen). Images were captured using a LSM710 confocal microscope (Carl Zeiss, Germany). Image processing and analysis were performed using Image J software. The V5 mean nuclear intensity signal was determined using the Image J plugin/algorithm, measured as previously ^7^. A minimum of 100 cells were included per condition.

Immunohistochemistry and immunofluorescence on xenograft sections Animal procedures involving HCT116 and OE21 xenografts were performed in accordance with current UK legislation and were approved by the University of Oxford Biomedical Services Ethical Review Committee, Oxford, UK. HCT116 and OE21 cells were grown as xenograft tumors as previously described ^52^. Briefly, 6–8-week-old female athymic nude mice (BALB/c nude) (Charles River, UK) were injected subcutaneously into the flank with 5 × 10^6^ HCT116 cells in serum-free DMEM. In addition, 6–8-week-old female CD-1 nude mice (Charles River, UK) were injected subcutaneously into the flank with 5 × 10^6^ OE21 cells in 50% (v/v) matrigel and serum-free RPMI. Two hours before the tumor was harvested, mice were injected intraperitoneally with 60 mg/kg of pimonidazole. Tumors were harvested before reaching 500 mm^3^ (according to the formula V=WxDxLxπ/6). Hypoxic regions were visualized by pimonidazole staining with hypoxyprobe 1 antibody (clone 4.3.11.3, Hypoxyprobe) after dewaxing and antigen retrieval with 10 mM sodium citrate buffer (pH 6.0). In addition, serial sections were stained for DDX5 (pAb204, Millipore), or no primary antibody (negative control), followed by HRP-conjugated secondary antibody incubation. Staining was developed with 3,3′-Diaminobenzidine (DAB, Vector Labs), and nuclei were counterstained with hematoxylin. Images were obtained using an Aperio Scanner (Leica Biosystems).

Orthotopic mouse GL261 glioma growth was conducted under the protocol 1019/2020 approved by the Local Ethics Committee for Animal Experimentation at the Nencki Institute of Experimental Biology in Warsaw. Male C57BL/6 mice (12 weeks) were anesthetized with 4% isofluorane and maintained at 1.5% isofluorane in oxygen during the tumor implantation procedure. A total of 8 ×10^4^ luc+tdT+ GL261 glioma cells in 1 µl of DMEM were inoculated into a right striatum at the rate of 0.25 μL per minute using a stereotactic apparatus. The coordinates for injection were +1 mm anterior-posterior, −2 mm lateral and −3 mm dorsal-ventral in relation to bregma. After 25 days post implantation animals were intraperitoneally injected with 60 mg/kg of pimonidazole and 2 h later were anesthetized, sacrificed and perfused with PBS and subsequently with 4% paraformaldehyde (PFA) in PBS. Brains with tumors were then removed and fixed additionally with 4% PFA in PBS for 24 h, followed by immersion in 30% sucrose for 48 h and mounting in OCT freezing medium on dry ice. Coronal sections 10 μm in size were collected using a cryostat. Sections were subjected to an antigen retrieval with 10 mM sodium citrate buffer (pH 6.0) and co-stained with hypoxyprobe 1 and DDX5 (ab126730, Abcam) antibodies was carried out. Nuclei were counterstained with DAPI. Images were acquired using a Zeiss LSM800 confocal microscope.

### qPCR

RNA samples were prepared using TRIzol (Invitrogen/Life Technologies). cDNA was synthesized from total RNA using the Verso Kit (Thermo Scientific). qPCR was performed with the SYBR Green PCR Master Mix Kit (Applied Biosystems) in a 7500 FAST Real-Time PCR thermocycler with v2.0.5 software (Applied Biosystems). Primer sequences are included in the supplementary information. Data were normalized to *18S* housekeeping gene. mRNA fold change was calculated using the 2–ΔΔCt method. The qPCR graphs show the mean ± SEM of three biological replicates and statistical significance was analyzed with two-tailed student t-tests for each gene.

### Gene expression correlations in TCGA datasets

RNA-sequencing data (RNA Seq V2 RSEM) for 382 colorectal adenocarcinoma, 166 glioblastoma, 408 bladder urothelial carcinoma, 1100 breast invasive carcinoma, 501 lung squamous cell carcinomas, and 517 lung adenocarcinomas patient sample datasets were extracted from the TCGA project, which can be accessed through cBioportal (http://www.cbioportal.org/). To examine the correlation of *DDX5* against the hypoxia signature, raw data for each sequenced gene were rescaled to set the median equal to 1, and the hypoxia signature was determined by quantifying the median expression of genes from the hypoxic signature ^53^. Log10 conversion of the hypoxia signature was plotted against Log10 conversion of raw data for *DDX5* (also rescaled to set the median equal to 1).

Correlations and statistical significance were determined by calculating Spearman’s rho rank correlation coefficients and two-tailed P values using Hmisc package in R studio.

### IvyGAP human glioblastoma data

Expression of hypoxia-inducible genes (*VEGFA, SLC2A1, CA9, LDHA, BNIP3, BNIP3L* and *LDHA*) as well as candidate genes belonging to the GO Splicing pathway (GO:0008380), whose promoters had decreased ATACseq signal in hypoxic conditions in our GL261 dataset, has been tested in the transcriptional atlas of human glioblastoma, where samples for the RNAseq analysis had been collected via microdissection from anatomic structures within glioblastoma (Ivy Glioblastoma Atlas Project; http://glioblastoma.alleninstitute.org/) ^31^. Microvascular Proliferation as well as Pseudopalisading Cells Around Necrosis regions were taken into account, which should clearly distinguish between oxygenated and hypoxic environments. The presented heatmap based on the expression z-scores was generated with an online selection tool at http://glioblastoma.alleninstitute.org/.

### ATACseq sequencing and data processing

ATACseq was performed as previously described with some modifications ^22^. While still in hypoxic conditions, GL261 cells were fixed with 1% formaldehyde (Thermo Scientific) for 10 min and then quenched with 0.125 M glycine for 5 min at room temperature. After returning to normal conditions (21% O_2_), 25,000 cells were washed and subsequently lysed in cold lysis buffer (10 mM Tris-HCl, pH 7.4, 10 mM NaCl, 3 mM MgCl2 and 0.1% IGEPAL CA-630) for 5 min on ice. Cells were then centrifuged at 500xg for 8 min and cell pellets resuspended in transposition reaction according to the standard ATACseq protocol using 2.5 ul of Tn5 enzyme per 25 000 of cells ^54^. After transposition cells were centrifuged and reverse crosslinked (50 mM Tris-Cl, 1 mM EDTA, 1% SDS, 0.2 M NaCl, 200 μg/ml proteinase K) at 65°C with an overnight shaking at 1000 rpm. Zymo DNA Clean & Concentrator-5 columns (ZymoResearch) were used to purify DNA and the sequencing libraries were prepared according to the original ATACseq protocol ^54^. The ATACseq libraries were assessed for appropriate quality using a Bioanalyzer 2100 and subjected to paired-end sequencing (2×76 bp) using NovaSeq 6000 (Illumina) at the Laboratory of Sequencing (Nencki Institute). The quality of raw fastq data was assessed using FASTQC software (https://www.bioinformatics.babraham.ac.uk/projects/fastqc/). Reads were trimmed using Trimmomatic to remove adapter and transposase sequence with option “sliding window 5:20” and with a cut off from 5’ and 3’ end set to 3 and 15, respectively ^55^. Reads with a length of less than 50 bp were discarded. Then paired-end reads were aligned to mm10 using Bowtie2 aligner v2.2.5 with parameters set to –very sensitive and -X2000 ^56^. Duplicate reads were subsequently removed with Picard (http://picard.sourceforge.net) and reads mapping to the mitochondrial genome were also removed. Only properly paired reads with mapping quality score (MAPQ) > 30 were kept for downstream analysis using Samtools view with option -q 30 ^57^. Peaks were called using MACS2 v.2.1.1.2 with parameters set to -f BAPME -q 0.001 – nomodel –shift 0 ^57^. The obtained peaks were then filtered for peaks overlapping mm10 ENCODE blacklisted genomic regions. To visualize the overlap of open genomic regions (peaks) from different conditions, the makeVennDiagram function from the ChIPpeakAnno package was used with minoverlap parameter set to 200 ^58^. All sequencing track Fig.s were generated by IGV using a normalized bigWig file created with deepTools (bamCoverage) and normalized to genome coverage - RPGC ^59,60^. ATACseq data files are available at https://www.ncbi.nlm.nih.gov/geo under accession number GSE200757.

### Differential analysis of ATACseq peaks

Changes in chromatin accessibility were assessed using DESeq2 ^61^. To create a set of consensuses ATAC peaks for the read count matrix, the following steps were performed. First, peaks for all replicates in each condition were intersected and only peaks with overlap > 200 bp were included. Then obtained peaks from all conditions were merged with reduction of overlapping regions using the “reduce” function from the GenomicRanges package to generate consensus peak lists ^62^. Peaks with FDR < 0.05 and |log2 fold change| >= 0.6 were classified as significantly different. Alternative ATACseq differential accessibility analysis was performed with csaw R package using aforementioned ATAC peak sets with either a trimmed mean of M values (TMM) or non-linear loess-based normalization method ^28,63^.

### Annotation for differential ATACseq peaks to genomic features

The peaks were annotated to genomic features using ChIPseeker with promoter region defined as +/- 2,000 bp around the transcription start site (TSS) ^64^. Each peak was annotated to only one genomic feature according to the default annotation priority in ChIPseeker. To obtain all the genes which had peaks were assigned around their promoter region, a custom-written script in R was used based on biomaRt and GenomicRanges packages ^62,65^. Functional enrichment analysis of differentially accessible regions nearby genes was performed using Gene Ontology, (GO) KEGG and Reactome databases ^25–27^.

### Statistical analysis and data processing

Statistical analyses for qPCRs, ChIP-qPCR, immunofluorescent analysis of DDX5 expression, 5’EU incorporation, RPA foci measurement and R-loop quantification were performed using GraphPad Prism software (GraphPad Software Inc.). As indicated in the Fig. legends, statistical tests included two-tailed student t-tests ns = non-significant, \**p* ≤0.05, \*\**p* ≤ 0.01, \*\*\**p* ≤ 0.001, \*\*\*\**p* ≤ 0.0001.

## AUTHORS CONTRIBUTIONS

KBL co-conceived the project; conceived, designed, performed or supervised, interpreted the majority of the experiments; and wrote the manuscript. MD performed ATACseq experiments, analyses and validation; and contributed to the writing of the manuscript. HE performed R-loop assays. J(utta)M performed qPCR analyses. EB contributed to the writing of the manuscript. AJG provided feedback and supervision. J(akub)M assisted with the ATACseq analyses and advised on the manuscript. BK advised on the manuscript and facilitated ATACseq experiments. EMH co-conceived the project; conceived, designed, and interpreted the majority of the experiments; and wrote the manuscript. All authors commented on the manuscript.

## ACKNOWLEDGEMENTS

Thank you to Drs Monica Olcina and Tiffany Ma (University of Oxford) for their insightful feedback on our manuscript. KBL and EB were supported by a CRUK grant C5255/A23755 (awarded to EMH). KBL was also supported by CRUKDF 0715-KL grant (awarded to KBL and EMH). MD and KBL were supported by National Science Centre (Poland) grant 2019/33/B/NZ1/01556 (awarded to KBL). HE was supported by Medical Research Council - UKRI grant MC _UU_00001/8 (awarded to AJG). We acknowledge Ms Beata Kaza for preparing GL261 cryo-sections. We acknowledge Mr Paweł Segit, Mr Karol Jacek and Mr Adria-Jaume Roura Canalda for guidance on ATACseq data analysis. We acknowledge Ms Paulina Szadkowska, Dr Bartlomiej Gielniewski and Dr Bartosz Wojtas for sequencing the ATACseq samples.

The authors disclose no potential conflicts of interest.

