## Supplementary Figures and Methods for "Hypoxia-mediated regulation of DDX5 through decreased chromatin accessibility and post-translational targeting restricts R-loop accumulation"

### SUPPLEMENTARY INFORMATION

Supplementary Figure legends Page 2

Supplementary Table legends Page 6

Primer sequences Page 7

siRNA sequences Page 8

References Page 9

### SUPPLMENTARY FIGURE LEGENDS

**Supplementary Figure 1. Oxygen-dependent changes in the chromatin accessibility**

**A.** A representative image of immunofluorescent staining of GL261 mouse glioma xenografts showing a hypoxic area co-stained for pimonidazole (PIMO) and Glut-1. Scale bar = 20 μm.

**B.** GL261 cells were exposed to hypoxia (Hyp, 0.1% $O_2$) for the times indicated and western blotting was carried out for the histone modifications indicated.

**C.** GL261 cells were exposed to hypoxia (Hyp, 0.1% $O_2$) for 24 h followed by reoxygenation (to 21% $O_2$) for the indicated times following hypoxia. Western blotting was carried out for the antibodies shown (representative of three independent experiments).

**D.** Percentage of increased (up) and decreased (down) ATACseq peaks for genomic annotations in cells treated with <0.1% $O_2$ or 1% $O_2$ hypoxia in relation to normoxic control.

**E, F.** Comparison of significantly increased **E** and decreased **F** ATACseq peaks at <0.1% $O_2$ versus 1% $O_2$ hypoxia at distinct genomic regions shown as a ratio of percentage of specific peaks.

**G.** Venn diagram showing the number of shared and distinct ATACseq peaks being repressed at the promoters in GL261 cells treated either with 1% or <0.1% $O_2$.

**Supplementary Figure 2. ATACseq data analysis using alternative normalization methods**

**A.** Volcano plots showing differentially altered ATACseq peaks in 1% $O_2$ (left graph) or <0.1% $O_2$ (right graph) versus 21% $O_2$ generated with the R csaw tool using predefined MACS2 peak sets with TMM-based normalization method [1]. Statistically significant peaks were defined for FDR < 0.05 and |log2 fold change| ≥ 0.6.

**B.** Volcano plots showing differentially altered ATACseq peaks in 1% $O_2$ (left graph) or <0.1% $O_2$ (right graph) versus 21% $O_2$ generated with the R csaw tool using predefined MACS2 peak sets with non-linear loess-based normalization method [1]. Statistically significant peaks were defined for FDR < 0.05 and |log2 fold change| ≥ 0.6.

**C, D.** Percentage of increased (up) and decreased (down) ATACseq peaks for genomic annotations in cells treated with <0.1% $O_2$ or 1% $O_2$ hypoxia in relation to normoxic control based on the csaw-TMM (**C**) and csaw-loess (**D**) normalization methods from (A) and (B).

**E.** A density plot of differential fold change (FC) (log2 value) of ATACseq peaks at <0.1% $O_2$ in relation to 21% $O_2$ for the three different normalization approaches used, including DEseq2 (black line), csaw-TMM (navy blue line) and csaw-loess (red line). For the csaw-loess normalization method, the log2FC values are the smallest (for decreased ATACseq peaks) compared to other methods, which is reflected by the smaller number of genes falling within the significance threshold set for repressed promoters of genes from the RNA Splicing pathway (GO:0008380).

**F.** A heatmap showing selected genes from RNA Splicing Pathway (GO:0008380) with decreased ATACseq signals within their promotor region at <0.1% $O_2$ in relation to 21% $O_2$ for three different normalization approach (deseq2 – default normalization, csaw – tmm normalization, csaw – loess normalization). Significantly decreased region was defined as FDR < 0.05 and |log2 fold change| ≥ 0.6).

**Supplementary Figure 3. Hypoxia-dependent chromatin accessibility changes at R-loop interacting factors**

**A.** Volcano plot from Fig. 1H highlighting repressed promoters in R-loop-interacting helicases at severe hypoxia (<0.1% $O_2$). The graph showing genes with significantly increased and decreased ATACseq signals within their promoter regions at <0.1% $O_2$ in relation to 21% $O_2$ with green dots representing all significantly regulated peaks (FDR < 0.05 and |log2 fold change| ≥ 0.6). The dots marked in black highlight the top R-loop-interacting factors shortlisted from [2].

**B-D.** ATACseq peak profiles (from Integrative Genomics Viewer, IGV) showing repressed chromatin at the promoters of *Dhx9* (**B**), *Setx* (**C**) *and Ddx5* (**D**) in GL261 cells.

**Supplementary Figure 4. Correlation of DDX5 expression and hypoxia metagene in TCGA cancer patient samples**

**A-F.** Dot plots showing correlation of expression of DDX5 mRNA with hypoxia metagene signature from Buffa et al., in the indicated TCGA cancer patient cohorts: **A** Glioblastoma, **B** Breast invasive carcinoma, **C** Bladder urothelial carcinoma, **D** Colorectal adenocarcinoma, **E** Lung adenocarcinoma and **F** squamous cell carcinoma [3]. Spearman's rank correlation coefficient and P values are shown for the Log10 median expression of DDX5 and hypoxic signature.

**Supplementary Figure 5. Hypoxia-dependent DDX5 repression in multiple cell lines**

**A.** HCT116 cells were exposed to the hypoxic conditions indicated, or hypoxia mimetics $CoCl_2$ (150 μM) or DFO (100 μM) for the times indicated. Western blotting was then carried out using the antibodies shown.

**B.** U87 cells were exposed to hypoxia (1% or <0.1% O2) for the times indicated followed by western blotting for DDX5 as well as HIF-1α hypoxic marker and β-actin (loading control).

**C-F.** H1299 **(C)**, RT112 **(D)**, U2OS **(E)** and MRC-5 **(F)** cells were exposed to hypoxia (Hyp, <0.1% $O_2$) for the times indicated followed by western blotting for DDX5 as well as HIF-1α hypoxic marker and β-actin (loading control).

**G, H.** HCT116 or RKO cells were transfected with siRNA duplexes against DDX5, DDX17 or DDX5/DDX17 and 48 h later subjected to 16 h of hypoxia (<0.1% $O_2$). Control siRNA (Scr) or no siRNA (Mock) were used as controls. Western blotting was carried out with the antibodies as indicated.

**I, J.** HCT116 cells were treated with emetine (20 μM) in hypoxic (Hyp, <0.1% $O_2$) or normoxic (Norm, 21% $O_2$) conditions for the times indicated and analyzed with western blotting. Densitometry is shown in **J**.

**K.** HCT116 cells were exposed to hypoxia in the presence of a neddylation inhibitor, MLN4924 (2 μM), as indicated, and western blotting carried out. A representative blot of three independent experiments is shown with densitometry underneath.

**Supplementary Figure 6. The effects of rescue of DDX5 expression in hypoxia on transcription and R-loops in HCT116 cells**

**A.** HCT116 cells were transfected with mycDDX5 or myc-tag control vector and western blotting was carried out with the indicated antibodies.

B. HCT116 cells transfected in A were exposed to hypoxia (8 h) with 5'EU (0.5 mM) added for the final hour. Staining for 5'EU was then carried out. Representative images are shown (scale bar = 10 μm). 5'EU staining in red, DAPI (blue) shows the nucleus.

**C**. Nuclear Intensity of 5'EU staining in B was determined. Data represents the mean expression and SEM from three independent experiments. Statistical significance was calculated with an unpaired student t-test for each indicated condition (* $p<0.05$).

**D.** HCT116 cells were transfected with myc control or myc-DDX5 and 24 h later exposed to hypoxia (<0.1% $O_2$). Western blotting was carried out with the antibodies indicated.

**E.** HCT116 cells were co-transfected with myc-tagged DDX5 and DHX9 together with V5-RNase H1$^{D210N}$ and exposed to hypoxia (18 h), followed by western blotting for the indicated antibodies

**F.** HCT116 cells were transfected as in part E and V5 fluorescence was analyzed. Representative images are shown (scale bar = 10 μm). V5 fluorescence is shown in green, DAPI (blue) shows the nucleus.

**G**. Nuclear fluorescence intensity of V5 (R-loops) was determined. Data represents the mean expression and SEM from four independent experiments. Statistical significance was calculated with an unpaired student t-test for each indicated condition (* $p<0.05$).

### SUPPLEMENTARY TABLE LEGENDS

**Supplementary Table 1. ATACseq differential peak analysis**

Differential ATACseq peak analysis in GL261 cell exposed to 16 h of hypoxic conditions: 1% $O_2$ and <0.1% $O_2$, in relation to normoxic control (21% $O_2$). Separate tabs show increased or decreased peaks at each hypoxic condition, accordingly. Fold change and p values were calculated with DESeq2 tool. Only statistically significantly changed peaks are included in the table. The peaks were annotated to genomic features with ChIPseeker.

**Supplementary Table 2. ATACseq differential peak analysis of promoter regions**

Differentially regulated ATACseq peaks in GL261 cells at promoter regions are shown and were generated as in Supplementary Table 1. In order to include assignment of peaks to multiple gene promoters the BiomaRt R package was used, as described in methods.

**Supplementary Table 3. ATACseq KEGG, GO and REACTOME pathway analysis at promoters repressed under severe hypoxia**

KEGG, GO and REACTOME pathways with significantly decreased ATACseq peaks at gene promoters identified in Supplementary Table 2 in response to <0.1% $O_2$ hypoxia treatment.

**Supplementary Table 4. ATACseq - spliceosome regulation in hypoxia**

KEGG, GO and REACTOME results showing downregulation of ATACseq peaks at the promoters of spliceosome pathways at <0.1% $O_2$ and 1% $O_2$. The analysis shows smaller amount of spliceosome genes having repressed ATACseq peaks at 1% $O_2$ in comparison to 0.1% $O_2$, hence the analysis of the whole pathways with KEGG, GO and REACOTME shows statistically insignificant change in the whole pathway at 1% $O_2$. However, some of the genes from the spliceosome pathways were still significantly repressed at 1% $O_2$, although with the smaller fold change than in <0.1% O2, and these are also shown in the tabs for the regulation under 1% $O_2$.

**Supplementary Table 5. ATACseq - R-loop-interactome**

ATACseq peaks significantly decreased at promoter regions of R-loop associated helicases in GL261 cells exposed to <0.1% $O_2$ in relation to normoxic control (candidate helicases shortlisted from [2]).

**PRIMER SEQUENCES**

**Human gene primers**

18S F: GCCCGAAGCGTTTACTTTGA

18S R: TCCATTATTCCTAGCTGCGGTATC

DDX5 F: GTGTCATCGGTGTCCTTCCT

DDX5 R: TAGAAAAGCGTGCGACAAGT

VEGF F: CTACCTCCACCATGCCAAGT

VEGF R: CTCGATTGGATGGCAGTAGC

**Mouse gene primers**

Srsf1 F: GTGGTTGTCTCTGGACTGCC

Srsf1 R: GTTGCTTCTGCTACGGCTTC

Sfpq F: TGTCGGTTGTTTGTGGGGAA

Sfpq r: GTGTGTGGCAAATCGAACCC

Alyref2 F: GTTTTCCTGGGTGCTGTTGTG

Alyref2 R: GTCATGTGTTCTGTCCATAAAAGT

Hnrnpu F: ACAACAGAGGTGGAATGCCC

Hnrnpu R: CCCTGCTGCCACTGATTGTA

Hnrnpk F: GTGCTGCCCTCACTCTACTG

Hnrnpk R: AGGTTGTGCACGTCCTTTGA

Srsf7 F: GATTGCAGGCAGAGGAGGTT

Srsf7 R: GTTTCTCCTCCATACCGCCC

Thoc1 F: CATTCTATTCTGCTGGCAAAAATTAT

Thoc1 R: AAAGAGTTGAATTCTTCCACAGAAAAC

Dhx9 F: ATACTTCCACGCCCTCATGC

Dhx9 R: AAGCCAAAACCACATCACGC

Dhx15 F: AGCAGCAATTCGGACAGTGA

Dhx15 R: CTGCTGGGGTGGAAGTGTAG

Ddx1 F: GGTGTCGACTGGAAAGCTCA

Ddx1 R: ATCTGGTTGTGCATCCGGTT

Vegfa F: GTCCGATTGAGACCCTGGTG

Vegfa R: GCTGGCTTTGGTGAGGTTTG

Glut1 F: ATCCCATCCACCACACTCAC

Glut1 R: GAGAAGCCCATAAGCACAGC

Rn18s F: CGGACATCTAAGGGCATCACA

Rn18s R: AACGAACGAGACTCTGGCATG

**SIRNA SEQUENCES**

siDDX5: AACUCUAAUGUGGAGUGCGAC

siDDX17: AACAAGGGUACCGCCUAUACC

siDDX5/17: GGCUAGAUGUGGAAGAUGU

*Supplementary Figure 1*

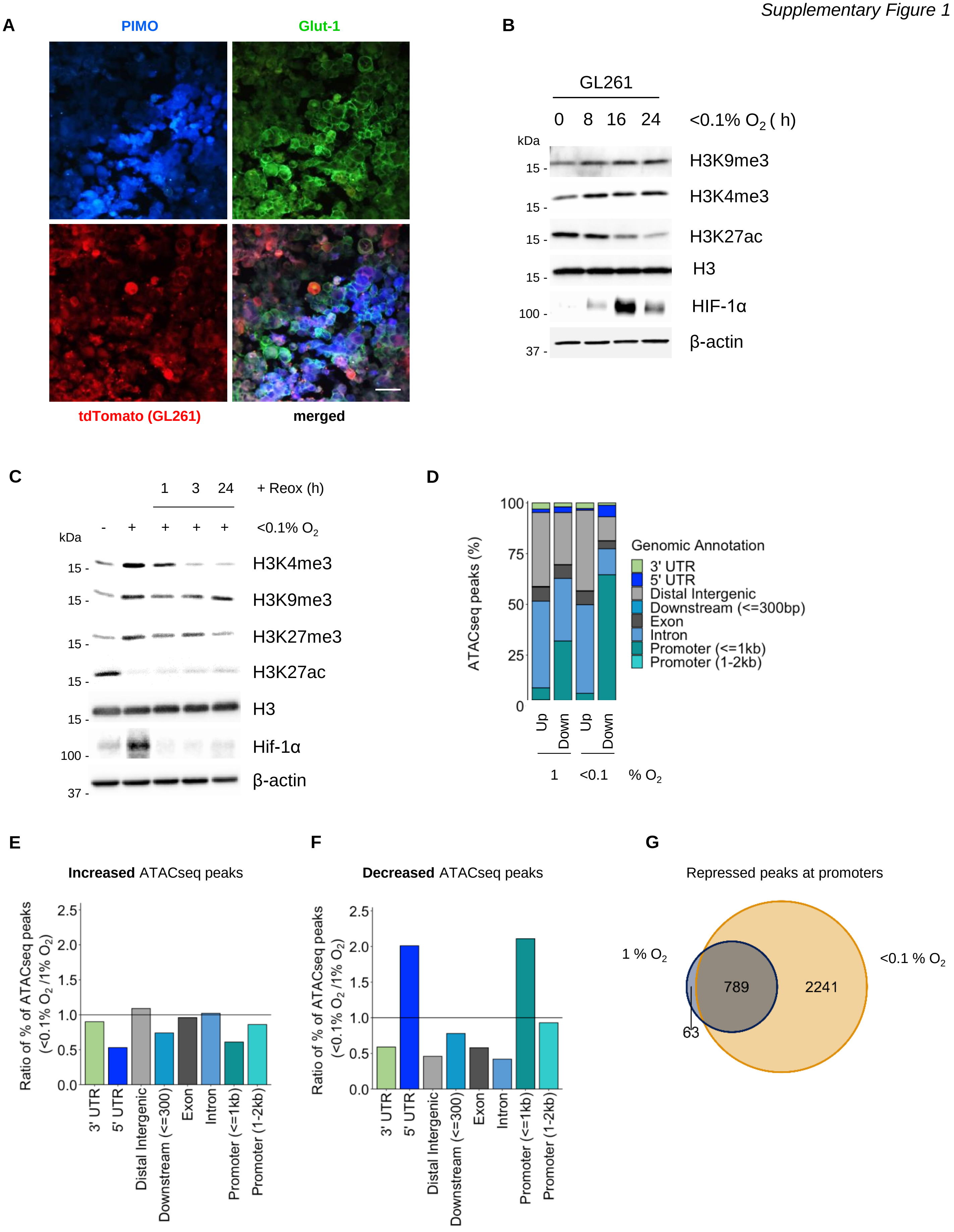

*Supplementary Figure 2*

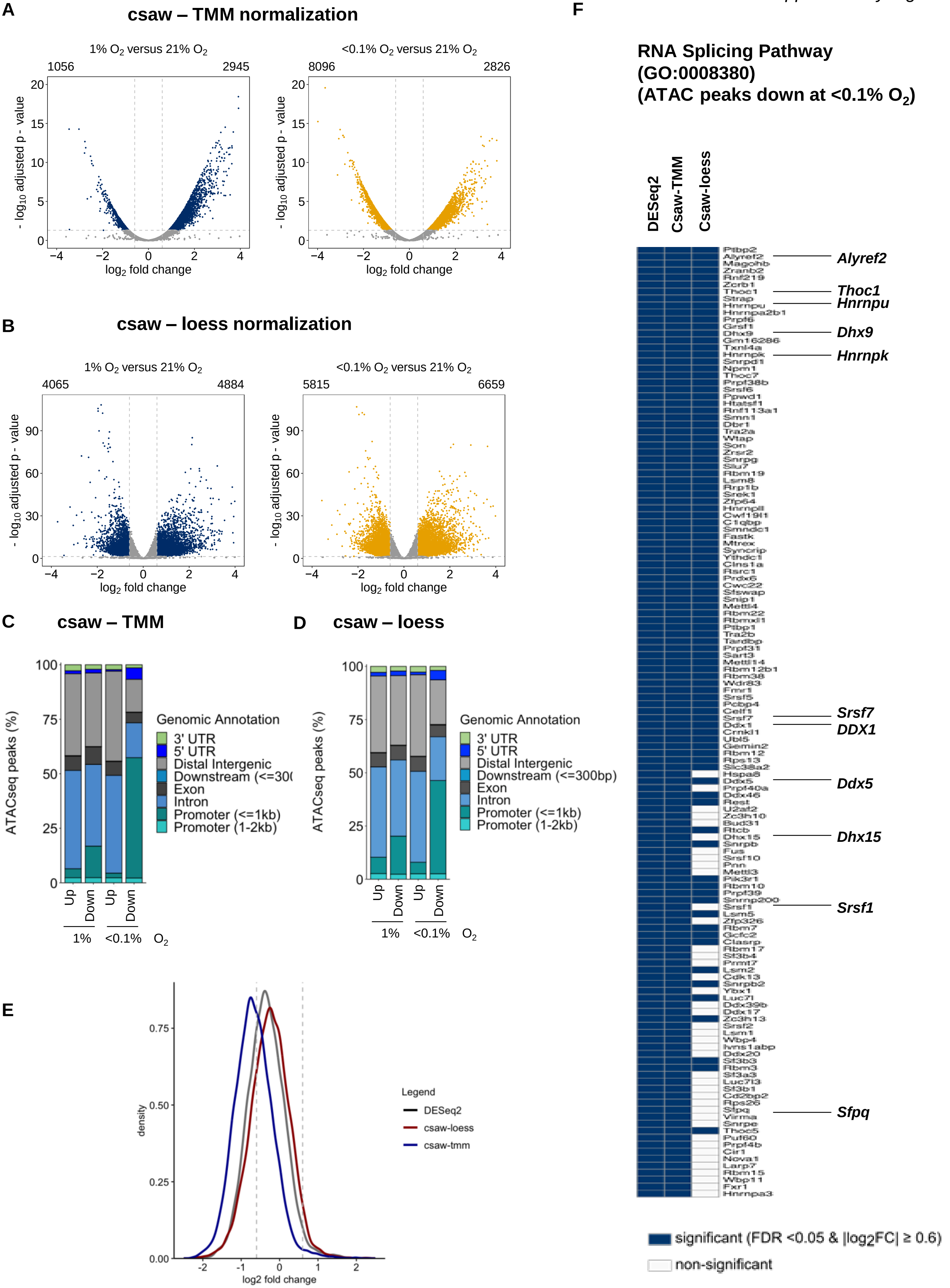

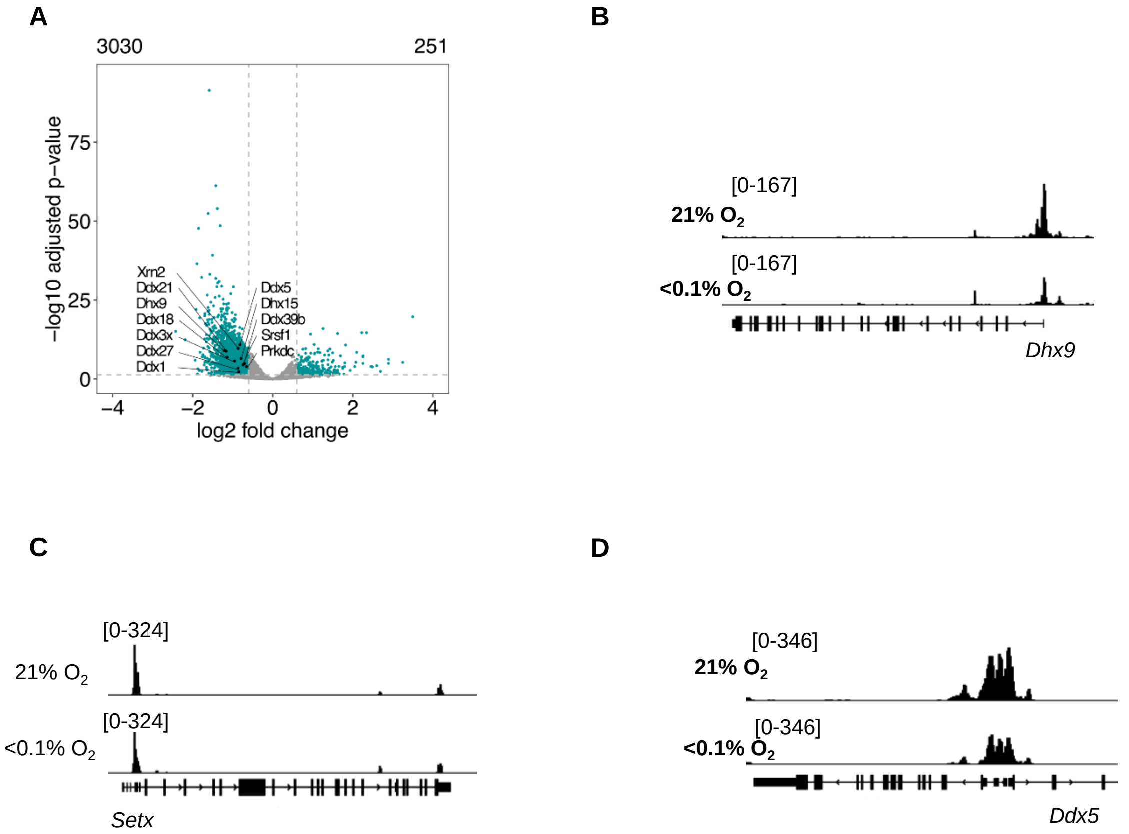
A
3030
251
−log10 adjusted p−value
75
50
25
0
−4
−2
0
2
4
log2 fold change
Xrn2
Ddx21
Dhx9
Ddx18
Ddx3x
Ddx27
Ddx1
Ddx5
Dhx15
Ddx39b
Srsf1
Prkdc
B
[0-167]
21% O2
[0-167]
<0.1% O2
Dhx9
C
[0-324]
21% O2
[0-324]
<0.1% O2
Setx
D
[0-346]
21% O2
[0-346]
<0.1% O2
Ddx5

*Supplementary Figure 4*

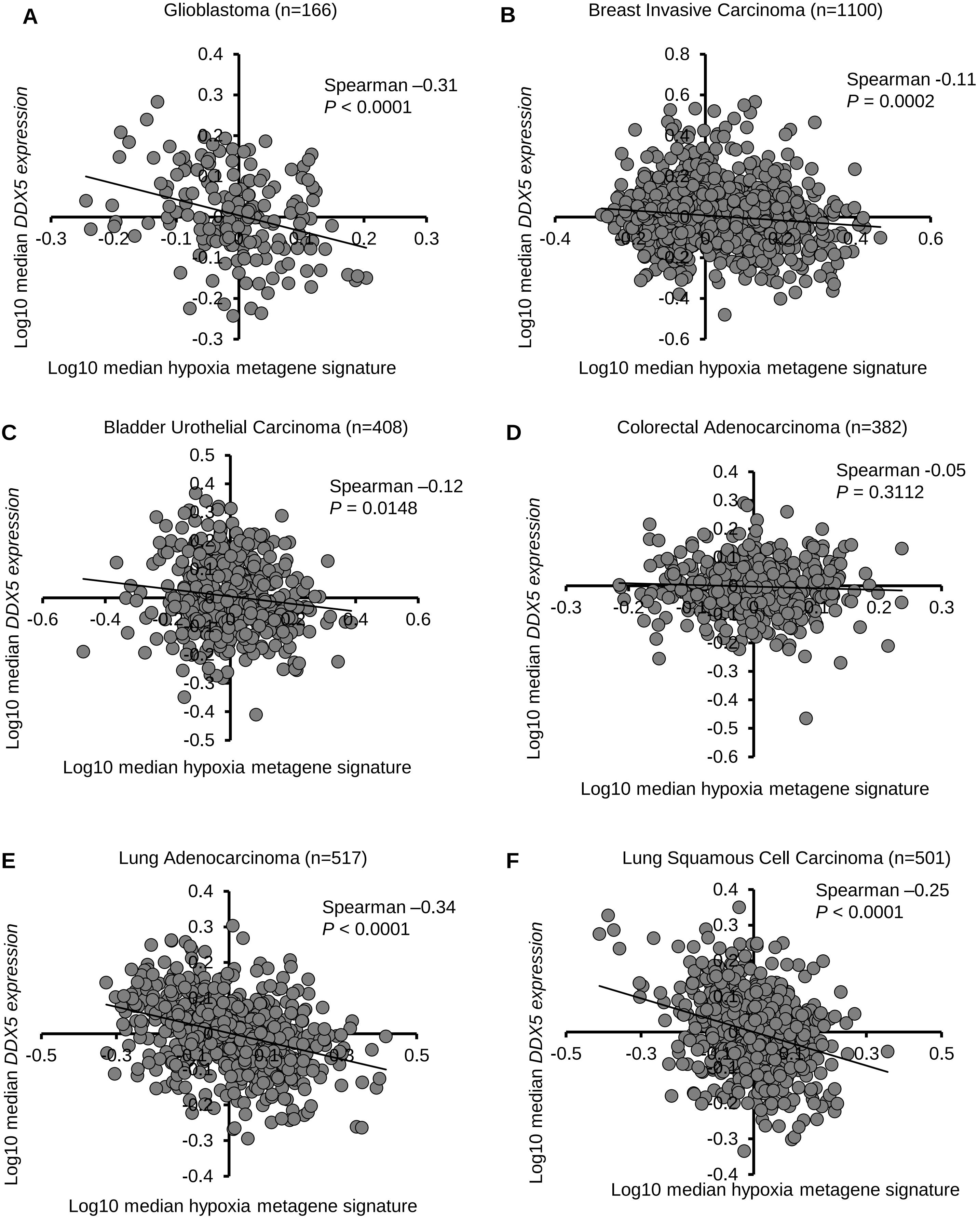

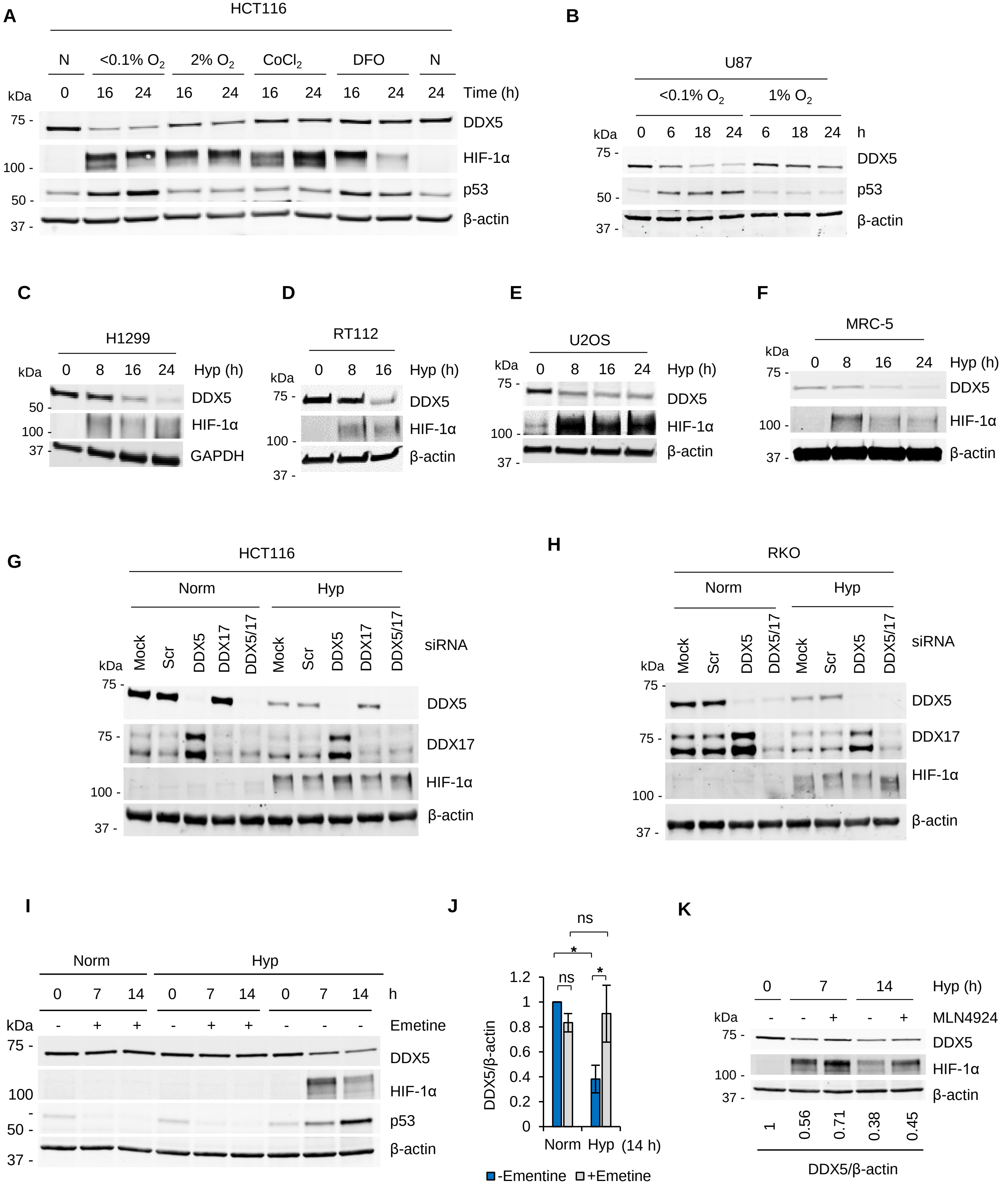
A
HCT116
N
<0.1% O2
2% O2
CoCl2
DFO
N
kDa
0 16 24 16 24 16 24 16 24 24 Time (h)
DDX5
HIF-1α
p53
β-actin
B
U87
<0.1% O2
1% O2
0 6 18 24 6 18 24 h
DDX5
p53
β-actin
C
H1299
0 8 16 24 Hyp (h)
DDX5
HIF-1α
GAPDH
D
RT112
0 8 16 Hyp (h)
DDX5
HIF-1α
β-actin
E
U2OS
0 8 16 24 Hyp (h)
DDX5
HIF-1α
β-actin
F
MRC-5
0 8 16 24 Hyp (h)
DDX5
HIF-1α
β-actin
G
HCT116
Norm
Hyp
Mock Scr DDX5 DDX17 DDX5/17 Mock Scr DDX5 DDX17 DDX5/17
siRNA
DDX5
DDX17
HIF-1α
β-actin
H
RKO
Norm
Hyp
Mock Scr DDX5 DDX5/17 Mock Scr DDX5 DDX5/17
siRNA
DDX5
DDX17
HIF-1α
β-actin
I
Norm
Hyp
0 7 14 0 7 14 0 7 14 h
- + + - + + - - - Emetine
DDX5
HIF-1α
p53
β-actin
J
DDX5/β-actin
Norm Hyp (14 h)
ns
*
-Ementine
+Emetine
K
0 7 14 Hyp (h)
- - + - + MLN4924
DDX5
HIF-1α
β-actin
1 0.56 0.71 0.38 0.45
DDX5/β-actin

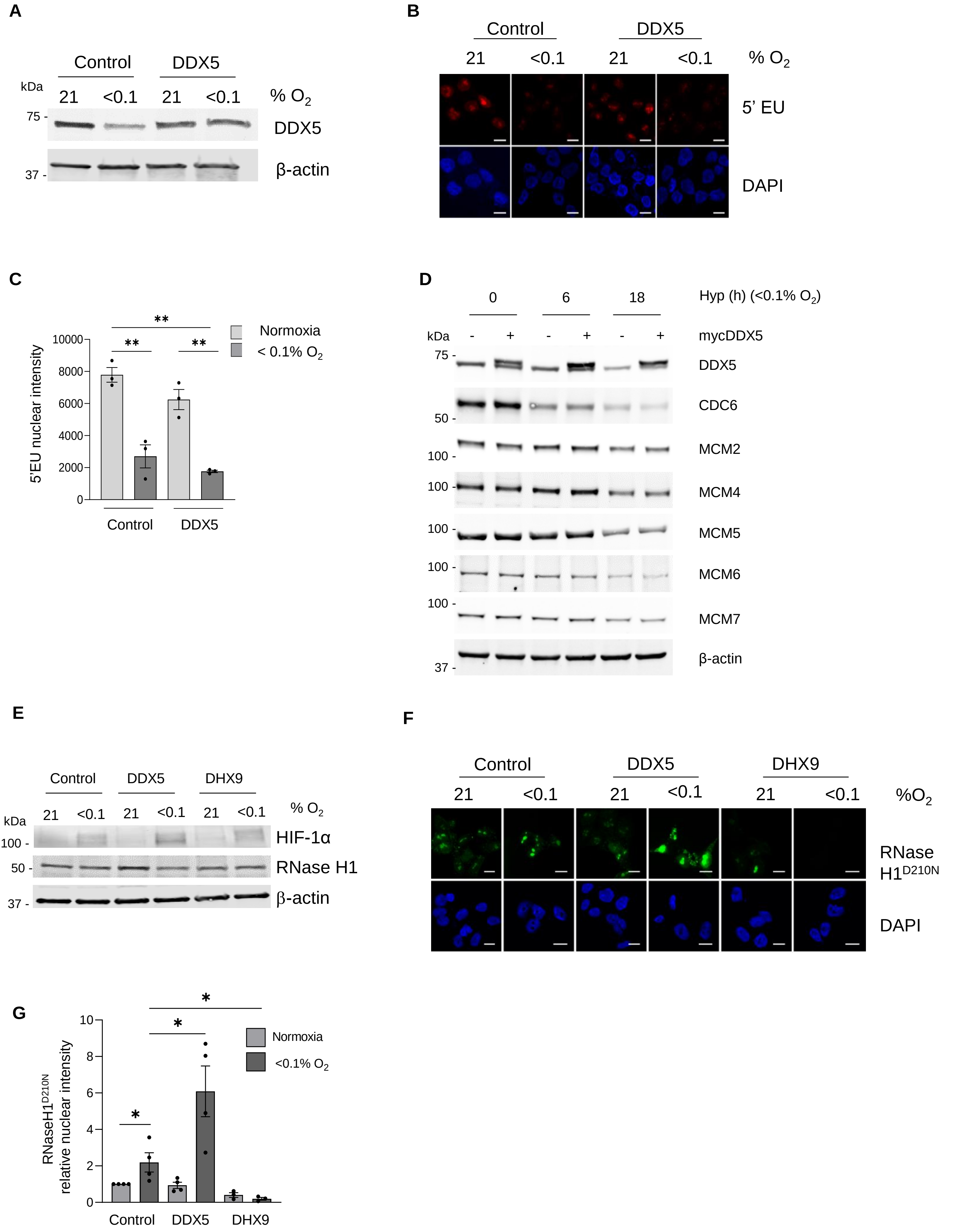
A
Control
DDX5
kDa
21
<0.1
21
<0.1
% O2
75 -
DDX5
37 -
β-actin
B
Control
DDX5
21
<0.1
21
<0.1
% O2
5' EU
DAPI
C
**
**
**
Normoxia
< 0.1% O2
5'EU nuclear intensity
10000
8000
6000
4000
2000
0
Control
DDX5
D
0
6
18
Hyp (h) (<0.1% O2)
kDa
-
+
-
+
-
+
mycDDX5
75 -
DDX5
CDC6
50 -
MCM2
100 -
100
MCM4
100
MCM5
100 -
MCM6
100 -
MCM7
β-actin
37 -
E
Control
DDX5
DHX9
% O2
kDa
21
<0.1
21
<0.1
21
<0.1
HIF-1α
100 -
50 -
RNase H1
37 -
β-actin
F
Control
DDX5
DHX9
21
<0.1
21
<0.1
21
<0.1
%O2
RNase H1D210N
DAPI
G
*
*
*
Normoxia
<0.1% O2
RNaseH1D210N relative nuclear intensity
10
8
6
4
2
0
Control
DDX5
DHX9
